# Species differences in the cerebellar distribution of six members of the Kv1 channel subfamily

**DOI:** 10.1101/2020.10.06.328237

**Authors:** Olimpia E. Curran, Andrew W. Hubball, Philip D. Minor, Charles H. Knowles, Joanne E. Martin

## Abstract

Comprehensive studies on the distribution of the Kv1 subfamily have been performed in rat (Chung et al., 2001) and gerbil (Chung et al., 2005), but not in mouse or human. We hypothesized that species differences may exist in the localization of these proteins. Two sets of polyclonal antibodies to Kv1.1-6 were used. Immunohistochemistry was performed on archived, formalin-fixed tissue from disease-free human, monkey and mouse cerebellum. Mouse staining corresponded to that described in rat and gerbil, with strong Kv1.1 and Kv1.2 immunoreactivities in the basket cell pinceau at the base of Purkinje cells. Kv1.3, Kv1.4, Kv1.5 and Kv1.6 were predominantly expressed in Purkinje cells. Human and monkey samples showed a similar pattern to mouse for Kv1.1, Kv1.2, Kv1.3 and Kv1.5. However, little or no Purkinje cell staining was seen in the primates with Kv1.4 and Kv1.6, and strong stellate cell expression was noted. All staining was abolished by cognate peptide blocking. Similar distributions were seen with both sets of antibodies. We conclude that there are marked species differences in the distribution of Kv1.4 and Kv1.6 between primates and rodents. Choosing appropriate animal models for studying physiological and disease processes may prove vital for translating research outcomes into clinical applications.

## Introduction

Potassium channels, a diverse family of membrane proteins, play a crucial role in regulating membrane potential and contribute to many vital cellular processes. Voltage-gated potassium (Kv) channels, in particular, underlie electrical impulse generation in muscle, endocrine and nerve cells. In neurons, Kv channels are essential for regulation of synaptic transmission and are key determinants of neuronal excitability. Action potential durations, firing frequencies and interspike intervals may be determined by the activity of Kv channels in neurons (Hille, 2001). The differential neuronal distribution of Kv channel subunits may be important for regulating presynaptic and postsynaptic membrane excitabilities.

Kv channels form an exceedingly diverse group of channels due to their molecular complexity as reflected by numerous gene families, multiple genes per family, heteromultimeric combination of subunits, auxiliary subunits, alternative splicing and post-translational modifications (Gutman et al., 2005). Molecular cloning of Kv channels from *Drosophila* has identified four distinct subfamilies of Kv channel genes, homologous to the *Drosophila* genes *Shaker, Shab, Shaw,* and *Shal* (Pongs, 1992). In total, twelve subfamilies (Kv1-12) of the principle pore-forming Kv α-subunit have been identified to date (Gutman et al., 2005). Members of the Kv1 channel subfamily are the mammalian homologues of *Drosophila* Shaker subunits. Each Kv1 channel family member consists of α- and β-subunits (Pongs, 1992). Structurally, four α-subunits assemble into a homo- or heterotetrameric (Sheng et al., 1993; Sheng et al., 1994; Wang et al., 1993) complex together with modifying β-subunits. The α-subunits are sufficient to form Kv channels, while the β-subunits have an auxiliary function in mediating rapid inactivation of Kv channels. Eight members of the Kv1 family have been identified up to now (Kv1.1-1.8) (Gutman et al., 2005), and the distribution of at least six members of this family (Kv1.1-1.6) has been extensively described in various regions of the brain (Chung et al., 2000; Guan et al., 2006; Tsaur et al., 1992; Wang et al., 1994).

The cerebellum is one of the brain areas where pattern of Kv1 channel subunits expression has been studied in some detail (Chung et al., 2001; Chung et al., 2005; McNamara et al., 1996; Sheng et al. 1994; Veh et al., 1995; Wang et al., 1993; Wang et al., 1994). *In situ* hybridization and immunocytochemistry studies have demonstrated that several Kv1 channel subunits show region- and cell type-specific expression (Sheng et al., 1992; Wang et al., 1993). Two studies have previously described cerebellar distribution of the six members of the Kv1 subfamily channels (Kv1.1-1.6) in the rat (Chung et al., 2001) and gerbil (Chung et al., 2005). However, there have been no comprehensive reports on the localisation of these six proteins in the primates and mouse cerebellum. Thus, the aim of present study was to describe the cerebellar distribution of Kv1.1-1.6 channel subunits in mouse, monkey and human, and correlate it with known rodent expression. We hypothesized that species differences may exist in the cerebellar localization of these proteins. Our results indicate that there are marked differences in the cerebellar distribution of Kv1.4 and Kv1.6 between primates and rodents.

## Materials and methods

### Tissues used

Formalin-fixed paraffin-embedded, disease-free cerebellum tissue from mouse, monkey and human were used in this study and included: two strains of female mice (<6 month old) C57/Black6 (n=3) and CD1 (n=3), four specimens of cerebellar archival tissue from female Cynomolgus monkeys (< 2 years old) and three human adult cases without neurological or psychiatric disorders selected from the Royal London Hospital *post mortem* archives. Human brains were removed within 48 h after death. All animals were treated in accordance with the Animals (Scientific Procedures) Act 1986 and the procedures involving animals were approved by the Home Office in the United Kingdom. Ethical approval was obtained (Project 06/Q0501/52) from the Central Office for Research Ethics Committee for the use of human tissue blocks. The human tissue removed from the neurological disease-free cadavers was fixed in 10% formal saline for at least 24 hours prior to processing. Archival monkey tissue was kindly donated by Dr Philip Minor and was perfusion-fixed material embedded in paraffin and stored for at least 10 years prior to use. Monkeys had been killed by a lethal dose of intravenous anesthetic and perfusion-fixed with formal saline for unrelated studies prior to this work. The mice were culled by carbon dioxide intoxication and the cerebella removed following cervical dislocation. Mice cerebella were then fixed in 4% formal saline overnight and embedded into paraffin blocks according to standard histological procedures. Tissue slides were prepared with 3-μm-thick sections, which were then dewaxed, dehydrated and pre-incubated in 3% H_2_O_2_ (in methanol) to suppress endogenous peroxidase activity. Two sets of commercially available polyclonal antibodies to six Kv1.1-1.6 channel subunits were used in this study. Polyclonal anti-Kv1.1, Kv1.2, Kv1.3, Kv1.4, Kv1.5, and Kv1.6 (Alomone Labs, Jerusalem, Israel; and Santa Cruz Biotechnology, USA) were used as primary antibodies. All antibodies were used at optimized dilutions and incubation times (Table 1 and 2). Indirect immunohistochemistry using the avidin-biotin-peroxidase complex (ABC) method utilizing the Vectastain Elite ABC kit (Vector, Peterborough, UK) was carried out after incubation with primary antibodies and followed by DAB (3,3’-diaminobenzidine) development in order to visualize the reaction product. Sections were viewed using a Leitz light microscope and images processed using Adobe Photoshop 7.0 software. Visual assessment of staining was semi-quantitatively reported as strong (++), moderate (+), weak (+/-) or absent (-) by two independent observers.

**Table 1.** Alomone Labs (Jerusalem, Israel) antibody characterization and optimal dilutions used in the present study.

| ***Antigen*** | ***Immunogen*** | ***Species antibody***<br>***was raised in,***<br>***preparation, catalog***<br>***number*** | ***Dilution,***<br>***Incubation***<br>***time*** |
| --- | --- | --- | --- |
| **Anti-Kv1.1** | GST fusion protein<br>mapping to residues 416-495 of mouse Kv1.1 | Rabbit polyclonal,<br>#APC-009 | 1: 300,<br>Overnight |
| **Anti-Kv1.2** | GST fusion protein<br>mapping to residues 417-499 of rat Kv1.2 | Rabbit polyclonal,<br>#APC-010 | 1:200,<br>Overnight |
| **Anti-Kv1.3** | GST fusion protein<br>mapping to residues 471-523 of human Kv1.3 | Rabbit polyclonal,<br>#APC-002 | 1:70,<br>Overnight |
| **Anti-Kv1.4** | GST fusion protein<br>mapping to residues 589-655 of rat Kv1.4 | Rabbit polyclonal,<br>#APC-007 | 1:150,<br>Overnight |
| **Anti-Kv1.5** | GST fusion protein<br>mapping to residues 513-602 of mouse Kv1.5 | Rabbit polyclonal,<br>#APC-004 | 1:100,<br>Overnight |
| **Anti-Kv1.6** | GST fusion protein<br>mapping to residues 463-530 of rat Kv1.6 | Rabbit polyclonal,<br>#APC-003 | 1:100,<br>Overnight |

Blocking serum: goat (1:100, Dakocytomation, UK). Secondary antibody: biotinylated goat anti-rabbit (1:300, Dakocytomation, UK).

**Table 2.** Santa Cruz Biotechnology (USA) antibody characterization and optimal dilutions used in the present study.

| ***Antigen*** | ***Immunogen*** | ***Species antibody***<br>***was raised in,***<br>***preparation, catalog***<br>***number*** | ***Dilution,***<br>***Incubation***<br>***time*** |
| --- | --- | --- | --- |
| **Anti-Kv1.1** | Peptide mapping at the N-terminus (N-16) of Kv1.1 of human origin | Goat polyclonal,<br># sc-11182 | 1: 75,<br>60 min |
| **Anti-Kv1.2** | Peptide mapping at the N-terminus (N-16) of Kv1.2 of human origin | Goat polyclonal,<br># sc-11186 | 1:40,<br>60 min |
| **Anti-Kv1.3** | Peptide mapping within an internal region (N-20) of Kv1.3 of human origin | Goat polyclonal,<br># sc-17239 | 1:40,<br>Overnight |
| **Anti-Kv1.4** | Peptide mapping within an internal region (F-20) of Kv1.4 of human origin | Goat polyclonal,<br># sc-16179 | 1:100,<br>60 min |
| **Anti-Kv1.5** | Peptide mapping near the N-terminus (N-16) of Kv1.5 of human origin | Goat polyclonal,<br># sc-11671 | 1:50,<br>60 min |
| **Anti-Kv1.6** | Peptide mapping near the C-terminus (C-15) of Kv1.6 of human origin | Goat polyclonal,<br># sc-16183 | 1:40,<br>60 min |

Blocking serum: rabbit (1:100, Dakocytomation, UK). Secondary antibody: biotinylated rabbit anti-goat (1:300, Dakocytomation, UK).

### Antibody characterization

Summary of Alomone Labs antibodies characterization is presented in Table 1. Briefly, each antibody was raised in rabbits using GST fusion protein as the immunogen, affinity purified and examined for specificity on immunoblots of rat brain membranes. Anti-Kv1.1, Kv1.2, Kv1.3, Kv1.4, Kv1.5 and Kv1.6 antibodies were made against proteins listed below: Kv1.1, residues 416-495, HRETE GEEQA QLLHV SSPNL ASDSD LSRRS SSTIS KSEYM EIEED MNNSI AHYRQ ANIRT GNCTT ADQNC VNKSK LLTDV; Kv1.2, residues 417-499, YHRET EGEEQ AQYLQ VTSCP KIPSS PDLKK SRSAS TISKS DYMEI QEGVN NSNED FREEN LKTAN CTLAN TNYVN ITKML TDV; Kv1.3, residues 471-523, TLSKS EYMVI EEGGM NHSAF PQTPF KTGNS TATCT TNNNP NSCVN IKKIF TDV; Kv1.4, residues 589-655, PYLPS NLLKK FRSST SSSLG DKSEY LEMEE GVKES LCGKE EKCQG KGDDS ETDKN NCSNA KAVET DV; Kv1.5, residues 513-602, HRETD HEEQA ALKEE QGIQR RESGL DTGGQ RKVSC SKASF HKTGG PLEST DSIRR GSCPL EKCHL KAKSN VDLRR SLYAL CLDTS RETDL; Kv1.6, residues 463-530, NYFYH RETEQ EEQGQ YTHVT CGQPT PDLKA TDNGL GKPDF AEASR ERRSS YLPTP HRAYA EKRML TEV (manufacturer’s datasheet). Staining specificity to six Kv channels in the mouse, monkey and human brains was already confirmed by immunohistochemistry on rodent brains. The antibodies used in this study had been shown to display a specific reaction to the α-subunits of Kv1 channels in the rat and gerbil brain, including cerebellum (Chung et al., 2001; 2005). Negative controls using both primary antibody omission and peptide block preabsorption additionally verified staining specificity (Fig. 1G, H). Sections from these controls did not show any of the immunoreactivity described in this report.

**Figure 1.**
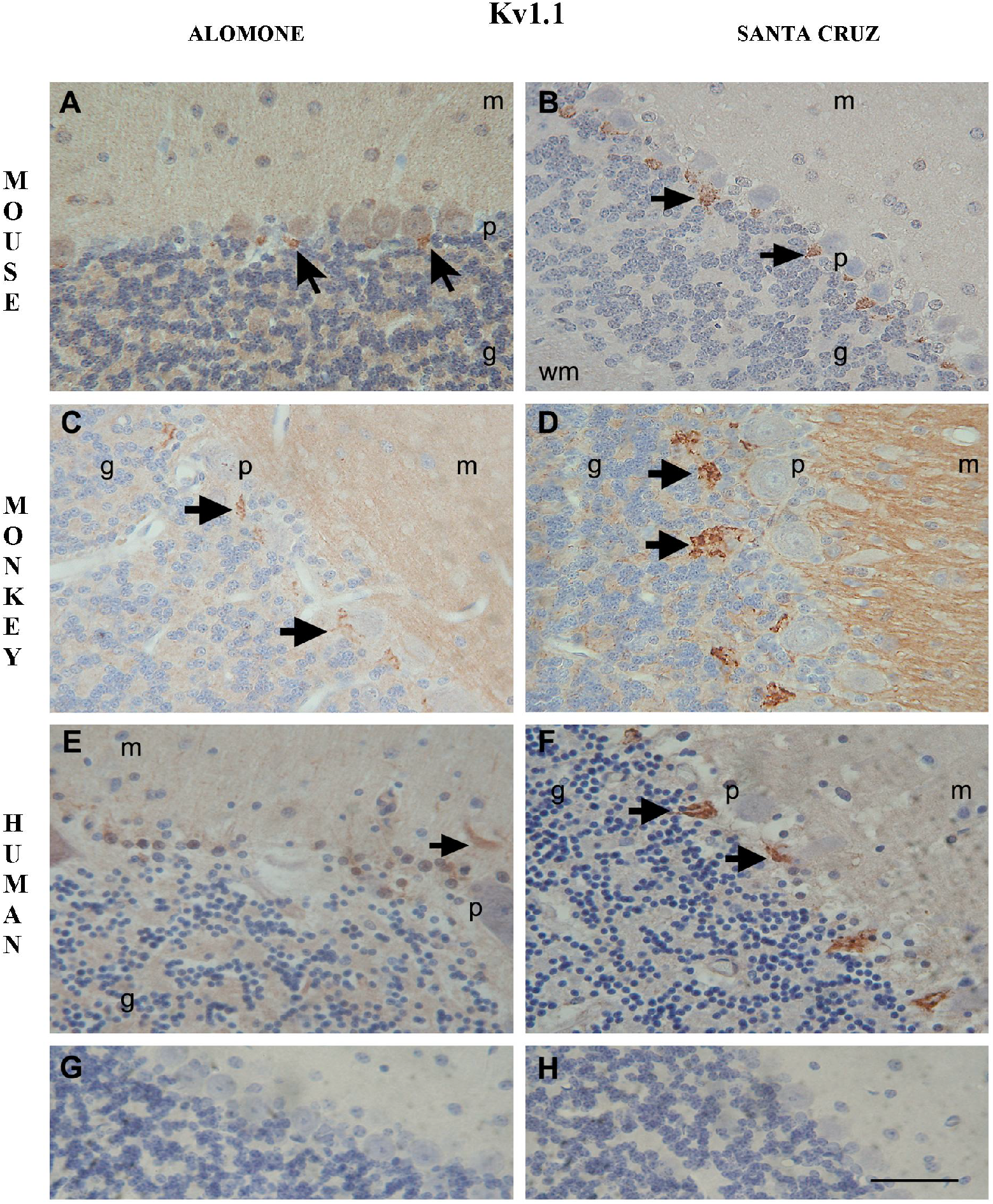
Cellular localizations of Kv1.1 subunit in the mouse (A, B); monkey (C, D) and human (E, F) cerebellar cortex. Immunohistochemistry was obtained using Alomone (A, C, E) and Santa Cruz (B, D, F) anti-Kv1.1 subunit antibodies. Immunoreactivities for Kv1.1 were concentrated in the basket cell axon plexus and terminal regions around the Purkinje cells (arrows). Low or absent immunoreactivity was detected in the Purkinje cell bodies. A sample of sections (shown here from mouse tissue) was reacted without primary antiserum (G), and a different sample was exposed to a primary antiserum that had been preincubated for 24 hours with control antigen peptides (H). No section from either group showed any of the immunoreactivity found in this report. m: molecular layer; p: Purkinje cell layer; g: granular layer; wm: white matter. All figures are based on following number of species: mouse (*n=6*), monkey (*n=4*), human (*n=3*); at least two sections per each animal were examined. Scale bar=50μm.

Summary of Santa Cruz Biotechnology antibodies characterization is presented in Table 2. Each antibody was raised in goat against an affinity purified peptide. Kv1.1, Kv1.2 and Kv1.3 showed a pattern of cellular morphology and distribution in the cerebellum that is identical to previous reports using corresponding Alomone Labs antibodies. Molecular weight of Kv1.1: 57-59 kDa, Kv1.2: 70 kDa and Kv1.4: 53 kDa. Kv1.4 antibody recognized a single band of 53 kD molecular weight on Western blots of SK-N-SH and U-87 MG whole cell lysates (manufacturer’s datasheet). Negative controls using both primary antibody omission and peptide block preabsorption verified staining specificity (Fig. 1G, H). Sections from these controls did not show any of the immunoreactivity described in this report.

## Results

Each of the six Kv1 α-channel subunits studied in this report had a unique pattern of distribution in the cerebellum (Table 3). As expected, extensive overlap in some areas of the cerebellum was also noted. No differences between our mouse strains were seen in any Kv1 localization study.

**Table 3.** Distribution of Kv1 α-channel subunits in mouse, monkey and human cerebellum.

| Mouse | Kv1.1 |  | Kv1.2 |  | Kv1.3 |  | Kv1.4 |  | Kv1.5 |  | Kv1.6 |  |
| --- | --- | --- | --- | --- | --- | --- | --- | --- | --- | --- | --- | --- |
|  | AL | SC | AL | SC | AL | SC | AL | SC | AL | SC | AL | SC |
| Molecular layer |  |  |  |  |  |  |  |  |  |  |  |  |
| Basket cells (BC) | +/- | - | +/- | +/- | +/- | +/- | + | + | +/- | +/- | + | + |
| BC endings | ++ | ++ | ++ | ++ | - | - | - | - | - | - | - | - |
| Stellate like cells | - | - | - | - | - | - | - | - | - | - | - | - |
| Purkinje cell layer |  |  |  |  |  |  |  |  |  |  |  |  |
| PC somata | +/- | - | +/- | +/- | + | + | ++ | ++ | ++ | ++ | ++ | ++ |
| Granular layer |  |  |  |  |  |  |  |  |  |  |  |  |
| Granule cells | - | - | - | - | - | - | - | - | - | - | - | - |
| Golgi cells | - | - | +/- | +/- | + | + | + | + | + | + | + | + |
| Cerebellar nuclei |  |  |  |  |  |  |  |  |  |  |  |  |
| Cell bodies | + | +/- | + | + | - | +/- | ++ | + | ++ | + | ++ | ++ |
| Neuropil | + | +/- | + | + | +/- | +/- | +/- | +/- | + | +/- | + | +/- |
| Stellate like cells | - | - | - | - | - | - | - | - | - | - | - | - |

| Monkey | Kv1.1 |  | Kv1.2 |  | Kv1.3 |  | Kv1.4 |  | Kv1.5 |  | Kv1.6 |  |
| --- | --- | --- | --- | --- | --- | --- | --- | --- | --- | --- | --- | --- |
|  | AL | SC | AL | SC | AL | SC | AL | SC | AL | SC | AL | SC |
| Molecular layer |  |  |  |  |  |  |  |  |  |  |  |  |
| Basket cells (BC) | - | - | +/- | +/- | +/- | +/- | - | - | +/- | - | +/- | - |
| BC endings | ++ | ++ | ++ | ++ | - | - | - | - | - | - | - | - |
| Stellate like cells | - | - | - | - | - | - | ++ | - | - | - | ++ | - |
| Purkinje cell layer |  |  |  |  |  |  |  |  |  |  |  |  |
| PC somata | - | - | +/- | +/- | + | + | - | - | ++ | - | +/- | - |
| Granular layer |  |  |  |  |  |  |  |  |  |  |  |  |
| Granule cells | - | - | - | - | - | - | - | - | - | - | - | - |
| Golgi cells | - | - | +/- | +/- | - | - | - | - | + | - | - | - |
| Cerebellar nuclei |  |  |  |  |  |  |  |  |  |  |  |  |
| Cell bodies | + | +/- | + | + | - | +/- | - | - | +/- | +/- | +/- | +/- |
| Neuropil | +/- | +/- | + | + | +/- | +/- | +/- | +/- | + | +/- | + | +/- |
| Stellate like cells | - | - | - | - | - | - | ++ | - | - | - | ++ | - |

| Human | Kv1.1 |  | Kv1.2 |  | Kv1.3 |  | Kv1.4 |  | Kv1.5 |  | Kv1.6 |  |
| --- | --- | --- | --- | --- | --- | --- | --- | --- | --- | --- | --- | --- |
|  | AL | SC | AL | SC | AL | SC | AL | SC | AL | SC | AL | SC |
| Molecular layer |  |  |  |  |  |  |  |  |  |  |  |  |
| Basket cells (BC) | - | - | +/- | +/- | +/- | +/- | - | - | +/- | - | +/- | +/- |
| BC endings | ++ | ++ | ++ | ++ | - | - | - | - | - | - | - | - |
| Stellate like cells | - | - | - | - | - | - | ++ | - | - | - | ++ | - |
| Purkinje cell layer |  |  |  |  |  |  |  |  |  |  |  |  |
| PC somata | - | - | +/- | +/- | + | + | - | - | + | +/- | +/- | +/- |
| Granular layer |  |  |  |  |  |  |  |  |  |  |  |  |
| Granule cells | - | - | - | - | - | - | - | - | - | - | - | - |
| Golgi cells | - | - | +/- | +/- | - | - | - | - | + | - | - | - |
| Cerebellar nuclei |  |  |  |  |  |  |  |  |  |  |  |  |
| Cell bodies | + | +/- | + | + | - | - | - | - | + | +/- | - | +/- |
| Neuropil | +/- | +/- | + | + | +/- | +/- | +/- | +/- | + | +/- | + | +/- |
| Stellate like cells | - | - | - | - | - | - | + | - | - | - | ++ | - |

Key: (++) strong staining, (+) moderate staining, (+/-) weak staining, (-) negative staining. AL – Alomone Labs antibody. SC – Santa Cruz Biotechnology antibody.

### Kv1.1

There was a high density of Kv1.1 in the cerebellum of mouse, monkey and human. Using both sets of Kv1.1 antibodies, immunoreactivity was found predominantly in the basket cell axon plexus and terminal regions around the Purkinje cells (Fig. 1). This pattern of staining appeared similar to that of Kv1.2 described below, but it was of relatively lower intensity. Purkinje cells appeared unstained as were granule cells of the granular layer. However, weak, patchy immunoreactivity was found in neuropil within the granular cell layer. In general, this distribution appeared the same for the three species investigated in this study and was in agreement with that previously described in mouse (Wang et al., 1994), rat (Chung et al., 2001) and gerbil (Chung et al., 2005). In the cerebellar nuclei, the cell bodies of cerebellar output neurons showed moderate staining for Kv1.1 using Alomone antibodies and a similar, but weaker pattern using Santa Cruz antibodies (Fig. 7).

### Kv1.2

Kv1.2 immunoreactivity was found in a number of different neuronal structures. In agreement with previous observations (McNamara et al., 1993; McNamara et al., 1996; Sheng et al., 1994; Wang et al., 1994), the most prominent staining was seen in the basket cell pinceau at the base of the Purkinje cell layer (Fig. 2). Lower intensity staining of some basket cell bodies in the molecular layer was also seen. In contrast to the basket cell pinceau, Purkinje cell bodies showed weak immunoreactivity for Kv1.2. Occasional staining of Golgi cells was detected in the granular layer, but granular cells themselves were unstained. As with Kv1.1, patchy immunostaining was also evident within the granular layer and weak punctate staining was present in the molecular cell layer. However, identity of these structures could not be ascertained at the light microscope level. The Kv1.2 distribution appeared grossly similar in the three species investigated in this study and was generally in agreement with that previously described in rat (Chung et al., 2001; McNamara et al., 1996; Sheng et al., 1994; Veh et al., 1995), mouse (Wang et al., 1994), and gerbil (Chung et al., 2005). The main discrepancy between species appears to be in the staining of Purkinje cell bodies with some researchers reporting some (Chung et al., 2005; Sheng et al., 1994), while others absent (Chung et al., 2001; McNamara et al., 1993; McNamara et al., 1996; Wang et al., 1994) Kv1.2 staining. Similarly to Kv1.1, immunoreactivity for Kv1.2 in the cerebellar nuclei was moderate in the cell bodies of cerebellar output neurons in all three species (Fig. 8).

**Figure 2.**
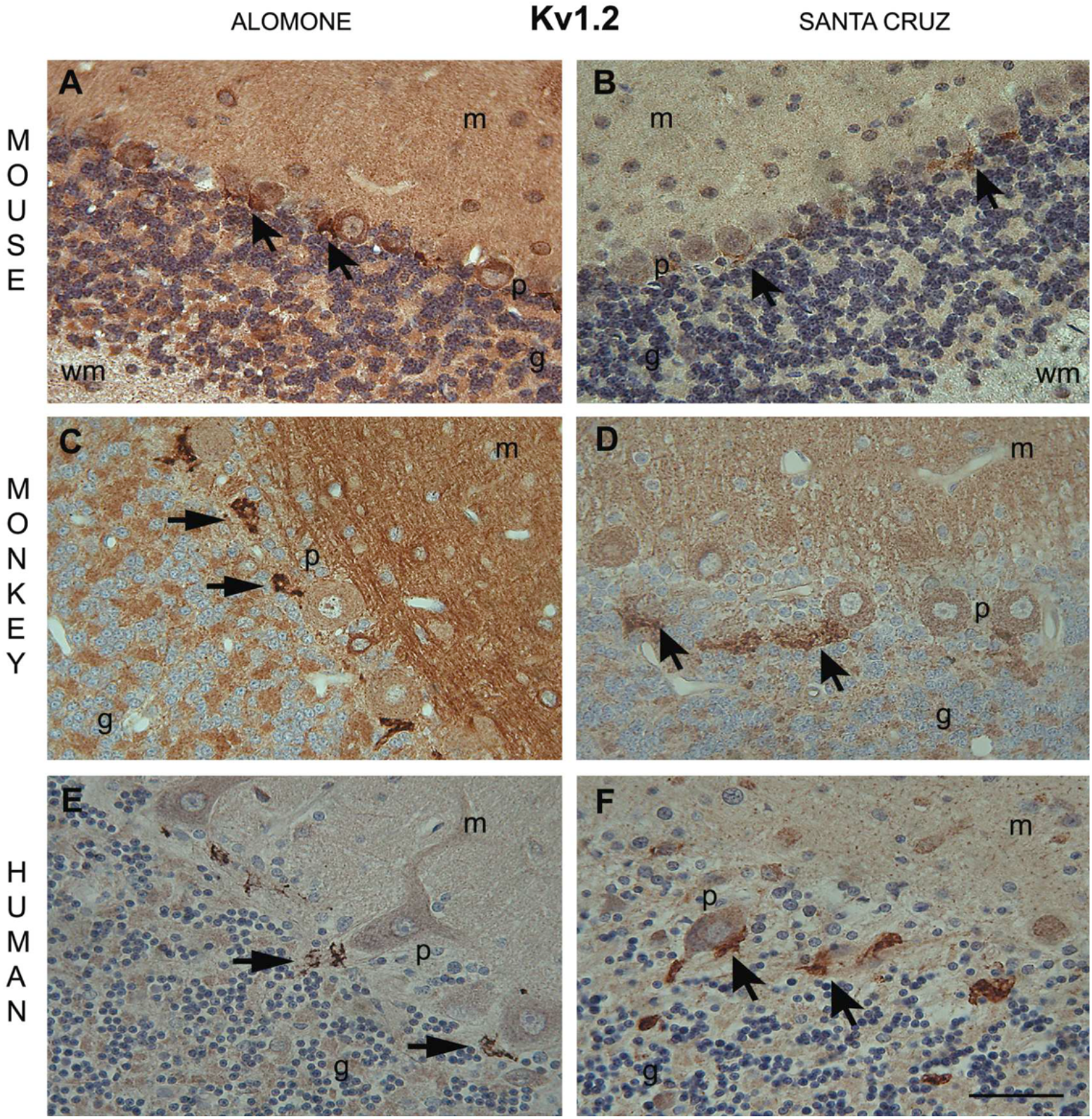
Cellular localizations of Kv1.2 subunit in the mouse (A, B); monkey (C, D) and human (E, F) cerebellar cortex. Immunohistochemistry was obtained using Alomone (A, C, E) and Santa Cruz (B, D, F) anti-Kv1.2 subunit antibodies. Similarly to Kv1.1, immunoreactivities for Kv1.2 were concentrated in the basket cell axon plexus and terminal regions around the Purkinje cells (arrows). Only low immunoreactivity was detected in the Purkinje cell bodies. m: molecular layer; p: Purkinje cell layer; g: granular layer; wm: whitematter. Scale bar=50μm.

### Kv1.3

In general, immunoreactivity for Kv1.3 was mostly confined to the somatodendric Purkinje cells (Fig. 3). There was weak immunoreactivity detected within the granular cell layer, although not in granule cells, in all three species. In addition, immunoreactivity was detected in basket cells of the molecular layer. Occasional Golgi cells were seen in the granular layer. This distribution is in agreement with previous reports in rat (Chung et al., 2001) and gerbil (Chung et al., 2005). Only weak staining of the cell bodies of cerebellar output neurons was detected for Kv1.3 in all three species (Fig. 9).

**Figure 3.**
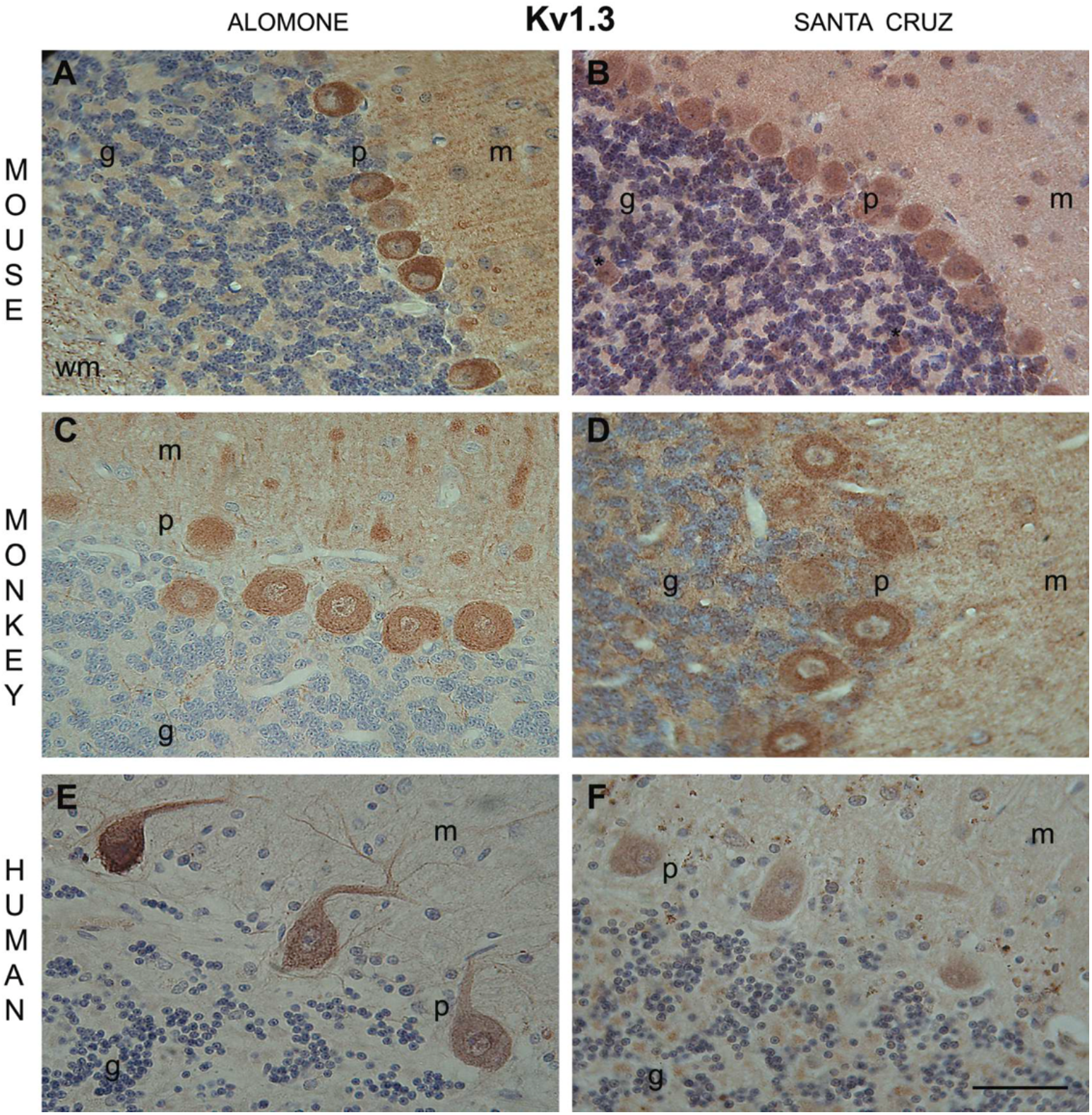
Cellular localizations of Kv1.3 subunit in the mouse (A, B); monkey (C, D) and human (E, F) cerebellar cortex. Immunohistochemistry was obtained using Alomone (A, C, E) and Santa Cruz (B, D, F) anti-Kv1.3 subunit antibodies. Immunoreactivities for Kv1.3 were found mostly in the somatodendritic Purkinje cells. m: molecular layer; p: Purkinje cell layer; g: granular layer; wm: white matter, * denotes Golgi type II. Scale bar=50μm.

### Kv1.4

In mouse, there was strong Kv1.4 immunoreactivity in Purkinje cell bodies with absent immunoreactivity in the granular layer, using both sets of antibodies (Fig. 4A, B). This pattern appeared similar to that reported in the gerbil (Chung et al., 2005). Some basket cells in the molecular layer and Golgi-like cells in the granular layer were also stained. In contrast, there was no immunoreactivity for Kv1.4 in Purkinje cells in monkey (Fig. 4C, D) and human (Fig. 4E, F) as previously described in the rat (Chung et al., 2001). This was true for both sets of antibodies. However, using Alomone antibodies only, in monkey and human, for the first time, we report a strong immunoreactivity in cells located in the molecular layer, which appeared to be stellate in form (Fig. 4C, E). Despite repeated runs of Santa Cruz antibodies, these cells were not seen in human and non-human primates. In the cerebellar nuclei, in mouse, generally strong Kv1.4 staining was detected in the cell bodies of cerebellar output neurons, using both sets of antibodies (Fig. 10A, B). In contrast, in primates, staining was absent from the soma of cerebellar output neurons (Fig. 10C, D, E, F). In addition, in primates, using Alomone antibodies, relatively strong (Fig. 10C, E) immunoreactivity was found in cells, which appeared to be stellate in form.

**Figure 4.**
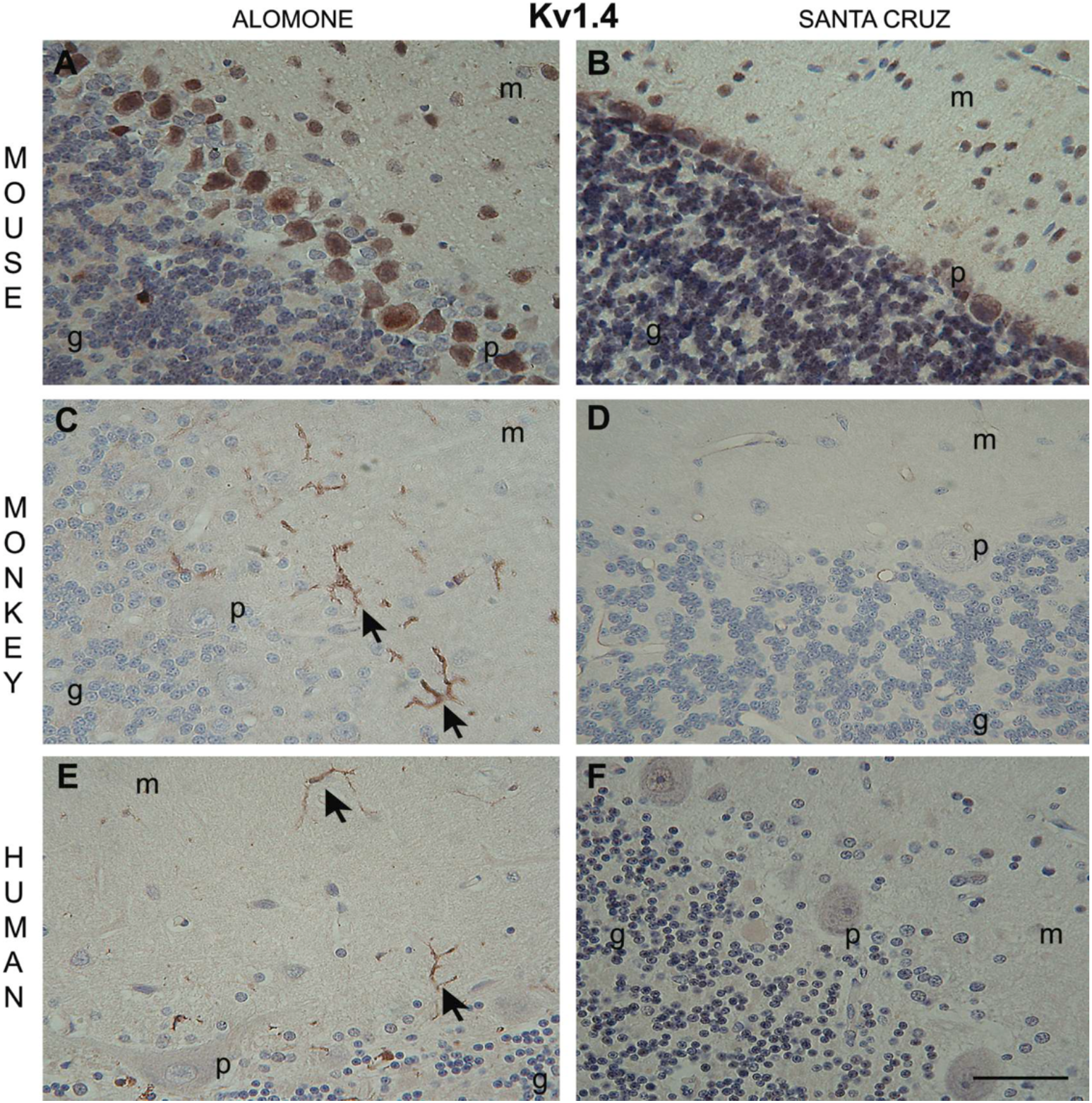
Cellular localizations of Kv1.4 subunit in the mouse (A, B); monkey (C, D) and human (E, F) cerebellar cortex. Immunohistochemistry was obtained using Alomone (A, C, E) and Santa Cruz (B, D, F) anti-Kv1.4 subunit antibodies. Strong Kv1.4 immunoreactivity was found in the Purkinje cell bodies in the mouse. In contrast, we observed no Purkinje cell immunoreactivity in monkey and human. In addition, in monkey and human tissue, using Alomone antibodies, for the first time, we report a strong immunoreactivity in cells in the molecular layer, which appear to be stellate in form (arrows). m: molecular layer; p: Purkinje cell layer; g: granular layer. Scale bar=50μm.

### Kv1.5

A high density of Kv1.5 was found in Purkinje cell bodies in all 3 species using both sets of antibodies (Fig. 5A, B, C, E). However, this pattern of staining in monkey and human was weaker using the Santa Cruz antibodies (Fig. 5D, F). In general, low intensity staining of Kv1.5 was also found in the molecular and granular layers. There was patchy, but significant signal detected in the granular layer, but granule cells appeared unstained. Occasional, but strong, immunoreactivity was present in large cells of the granular layer, consistent in appearance with Golgi cells. Grossly, Kv1.5 staining in mouse and primates was consistent with previous work in rodents (Chung et al., 2001; Chung et al., 2005). In the cerebellar nuclei, Kv1.5 proteins were clearly detected in the cell bodies of cerebellar output neurons in all three species, using Alomone antibodies (Fig. 11A, C, E). Similar, although weaker, cell body staining pattern was observed in mouse and monkey (Fig. 11B, D), and the weakest in human (Fig. 11F), using Santa Cruz antibodies.

**Figure 5.**
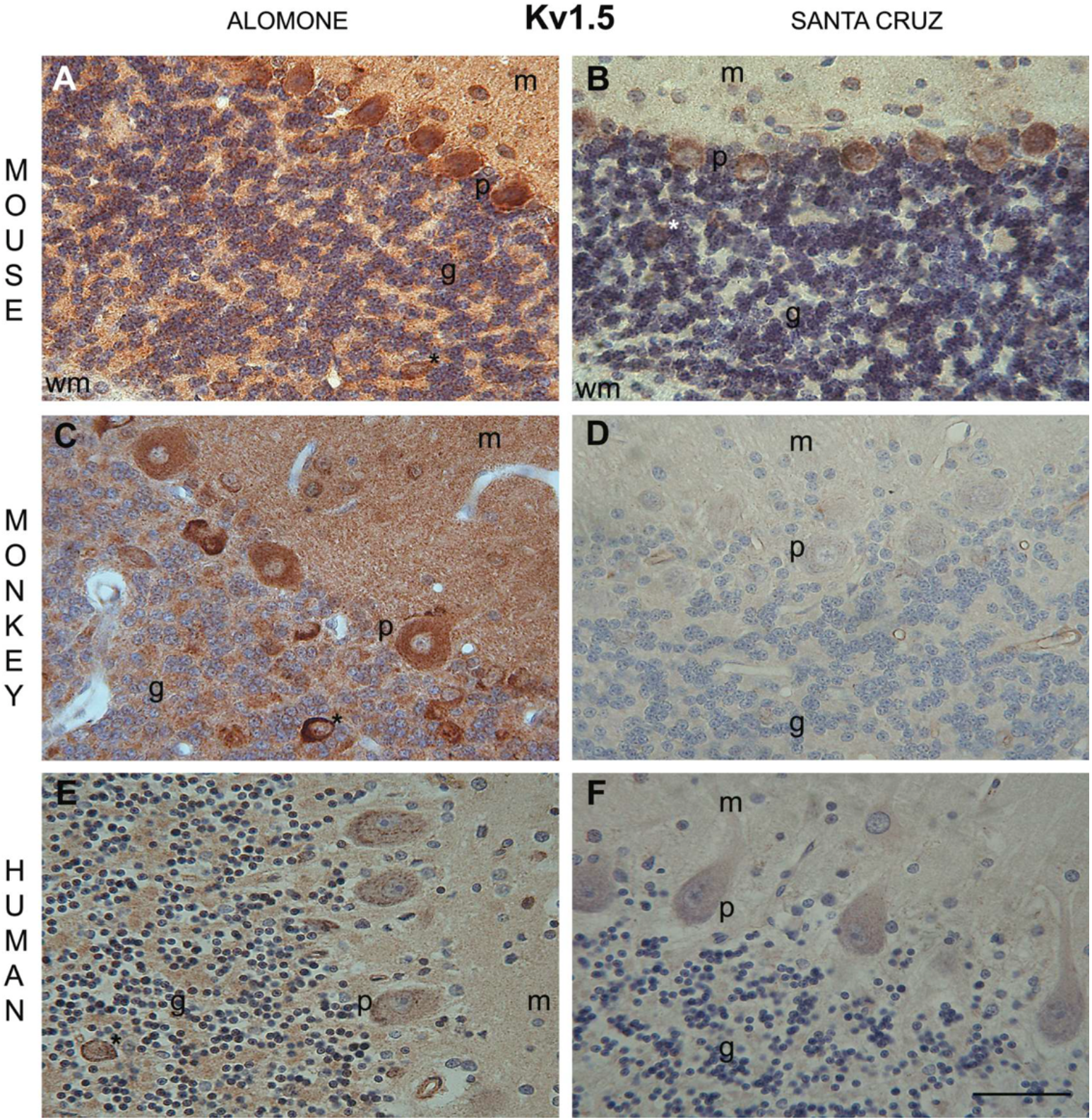
Cellular localizations of Kv1.5 subunit in the mouse (A, B); monkey (C, D) and human (E, F) cerebellar cortex. Immunohistochemistry was obtained using Alomone (A, C, E) and Santa Cruz (B, D, F) anti-Kv1.5 subunit antibodies. Kv1.5 immunoreactivity was predominantly found in the Purkinje cell bodies in mouse, using both antibodies, as well as in monkey and human, using Alomone antibodies. Weaker Purkinje cell bodies staining was noted in human and the weakest in monkey, using Santa Cruz antibodies. m: molecular layer; p: Purkinje cell layer; g: granular layer; * denotes Golgi cells type II. Scale bar=50μm.

### Kv1.6

In mouse, using both sets of antibodies, strong immunoreactivities for Kv1.6 were found in the Purkinje cell bodies, basket cells of the molecular layer, the granular layer and its Golgi-like cells. This is in keeping with reports in rat (Chung et al., 2001) and gerbil (Chung et al., 2005). The granule cells appeared unstained (Fig. 6A, B). In contrast, only very weak (Fig. 6E, F) or absent immunostaining was found in Purkinje cell bodies in primates (Fig. 6C, D). In addition, using Alomone antibodies only, we found relatively strong immunoreactivity in the molecular layer for what appeared to be stellate in form cells (Fig. 6C, E). In the cerebellar nuclei, Kv1.6 immunoreactivity was predominantly found in the cell bodies of cerebellar output neurons in mouse, using both antibodies (Fig. 12A, B). However, only weak or absent signal was found in cell bodies in primates (Fig.12C, D, E, F). In addition, in primates, using Alomone antibodies, immunoreactivity for what appear to be stellate in form cells was clearly detected (Fig. 12C, E).

**Figure 6.**
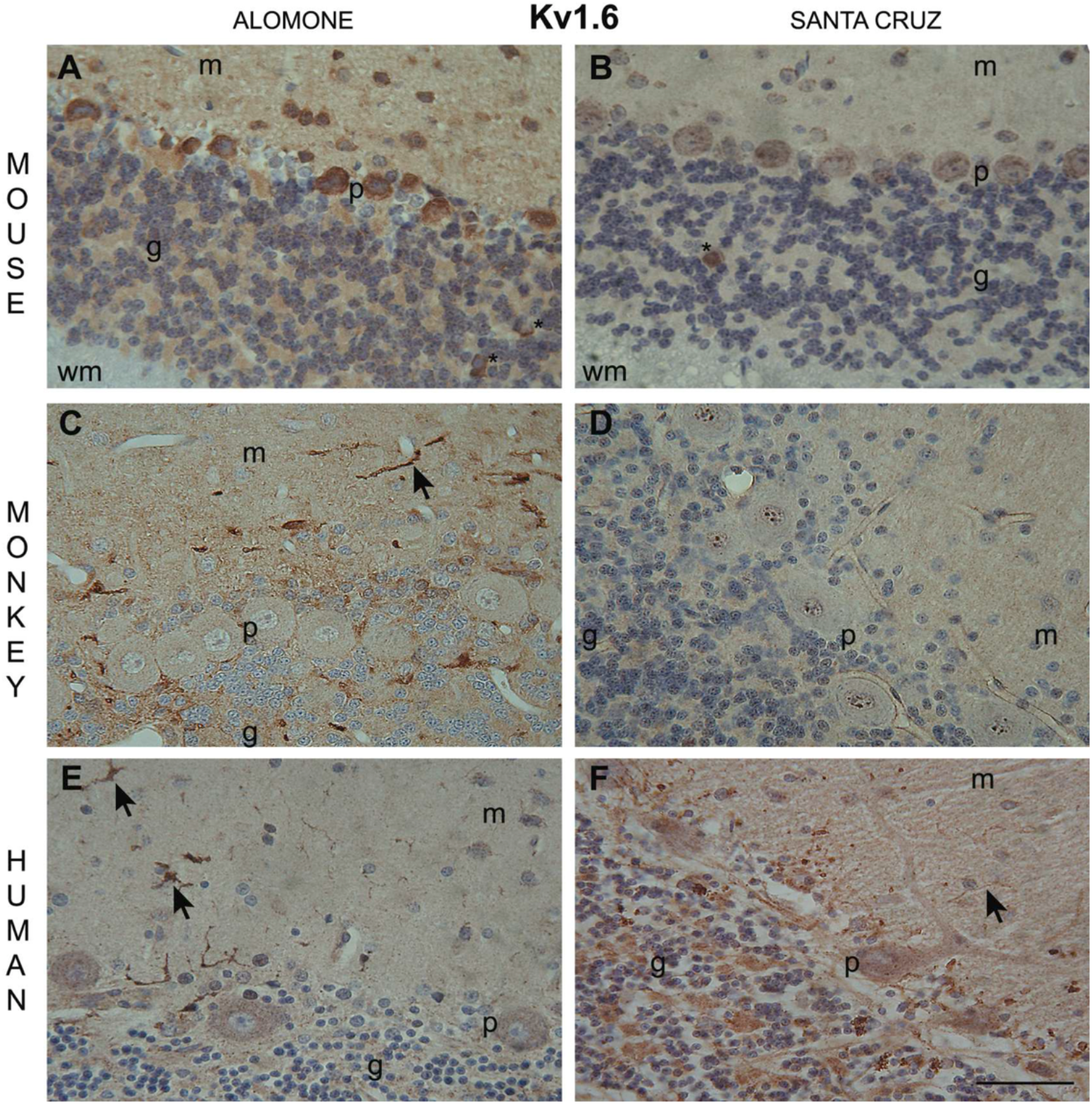
Cellular localizations of Kv1.6 subunit in the mouse (A, B); monkey (C, D) and human (E, F) cerebellar cortex. Immunohistochemistry was obtained using Alomone (A, C, E) and Santa Cruz (B, D, F) anti-Kv1.6 subunit antibodies. Kv1.6 immunoreactivity was predominantly found in the Purkinje cell bodies in mouse, using both antibodies. However, only weak or absent signal was found in Purkinje cell bodies in primates. In addition, using Alomone antibodies, we found relatively strong immunoreactivity for what appear to be stellate in form cells (arrows). m: molecular layer; p: Purkinje cell layer; g: granular layer; * denotes Golgi cells type II. Scale bar=50μm.

**Figure 7.**
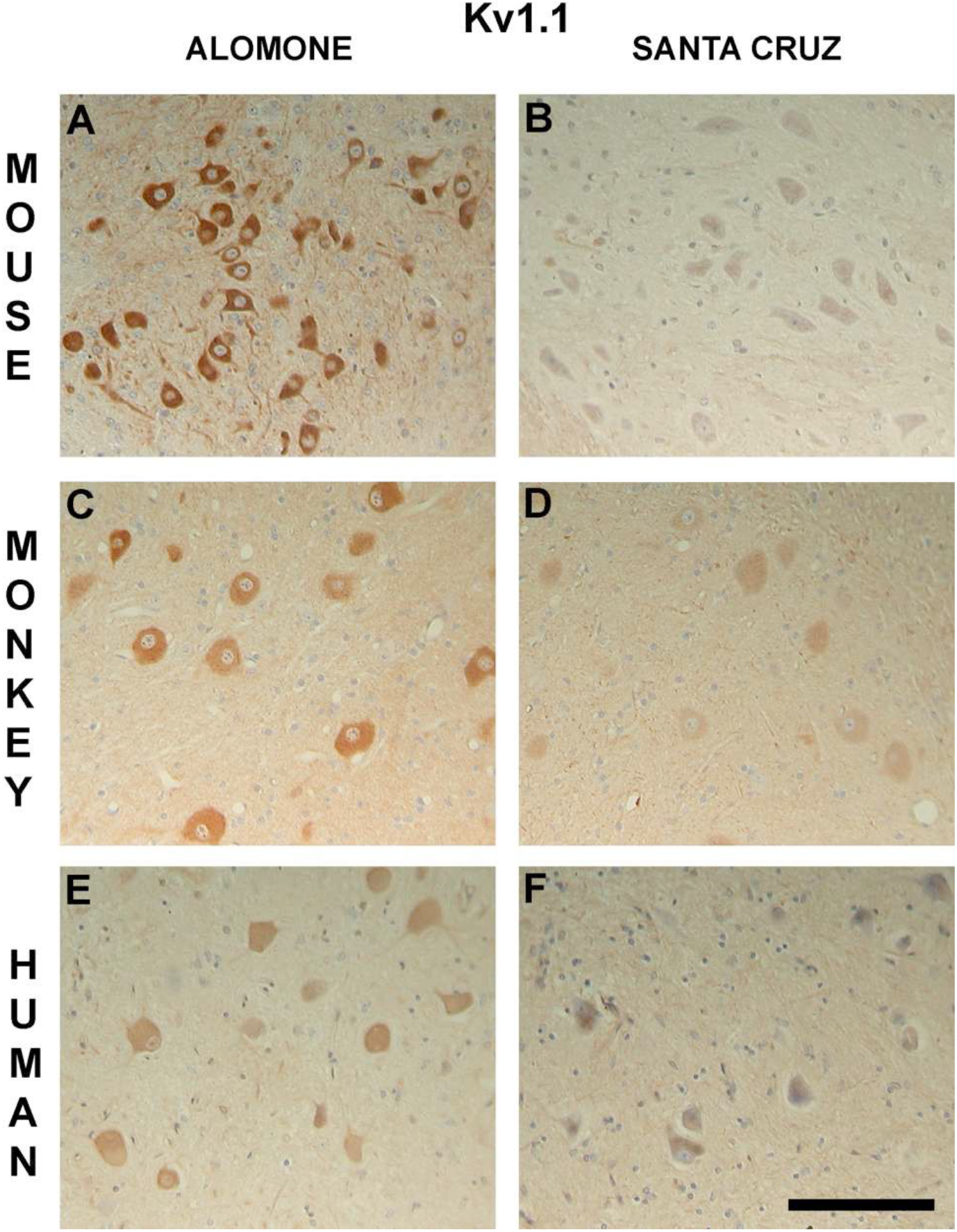
Cellular localizations of Kv1.1 subunit in the mouse (A, B); monkey (C, D) and human (E, F) cerebellar nuclei. Immunohistochemistry was obtained using Alomone (A, C, E) and Santa Cruz (B, D, F) anti-Kv1.1 subunit antibodies. Kv1.1 staining was clearly detected in the cell bodies of cerebellar output neurons, especially using Alomone antibodies (A, C, E). Weak staining was also detected in the surrounding neuropil. Similar, but weaker pattern was observed in tissues stained with Santa Cruz antibodies (B, D, F). Representative sections of the nucleus medialis (fastigial), interpositus and lateralis (dentate) in the mouse and monkey tissues and the nucleus lateralis (dentate) in the human tissue. All figures are based on following number of species: mouse (*n=6*), monkey (*n=4*), human (*n=3*); at least two sections per each animal were examined. Scale bar=125μm.

**Figure 8.**
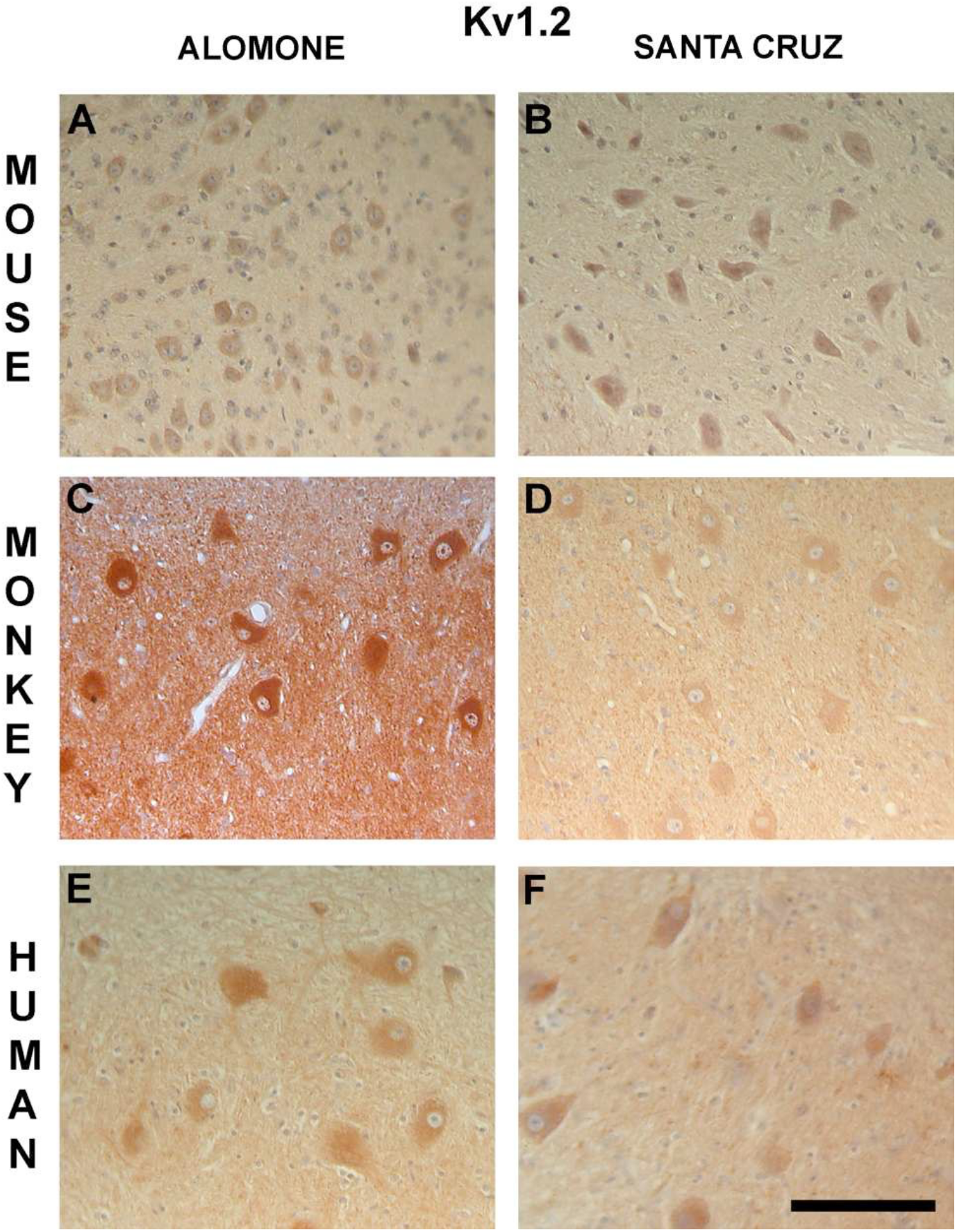
Cellular localizations of Kv1.2 subunit in the mouse (A, B); monkey (C, D) and human (E, F) cerebellar nuclei. Immunohistochemistry was obtained using Alomone (A, C, E) and Santa Cruz (B, D, F) anti-Kv1.2 subunit antibodies. Kv1.2 proteins were clearly detected in the soma of cerebellar output neurons. Weaker staining was observed in the surrounding neuropil. Scale bar=125μm.

**Figure 9.**
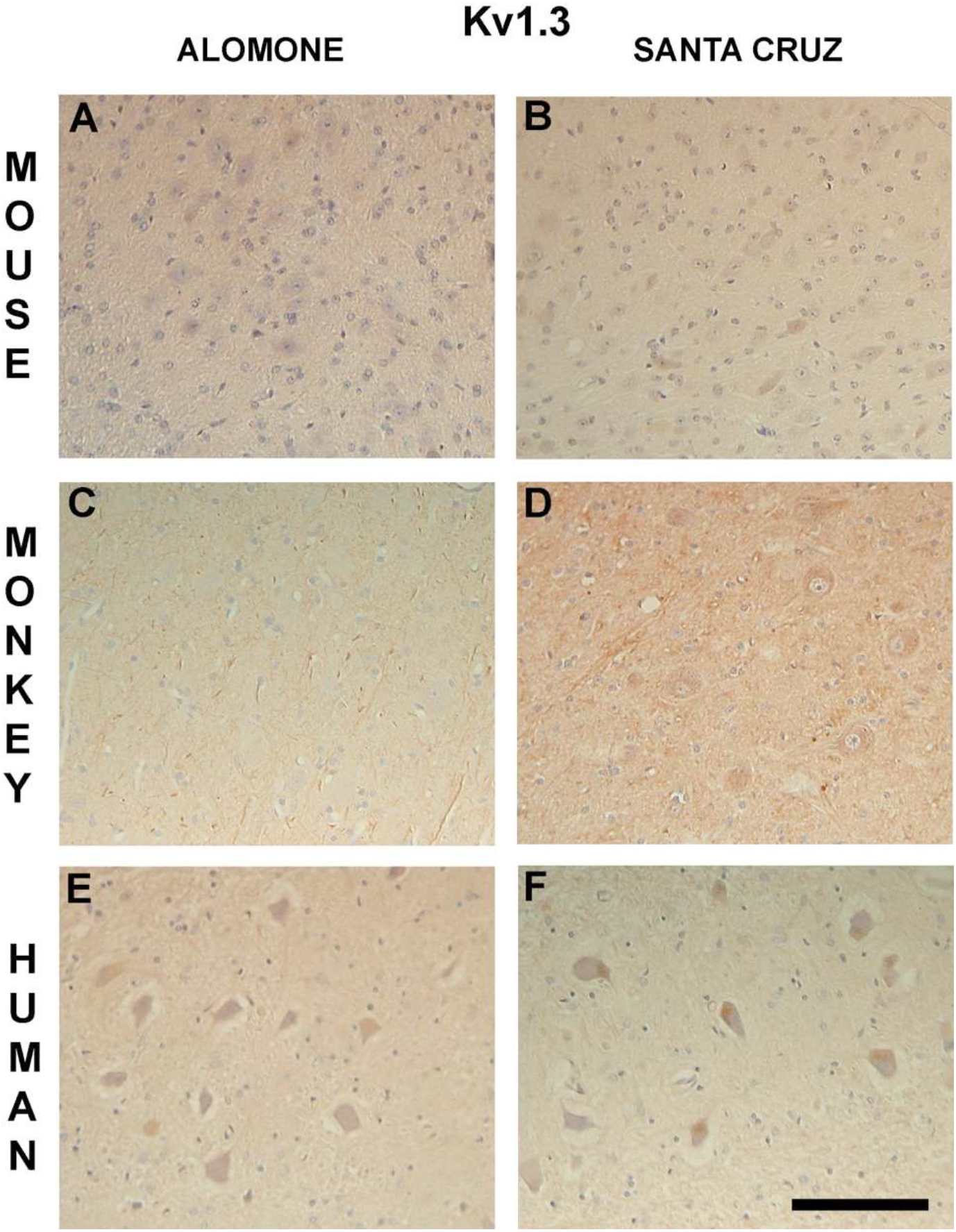
Cellular localizations of Kv1.3 subunit in the mouse (A, B); monkey (C, D) and human (E, F) cerebellar nuclei. Immunohistochemistry was obtained using Alomone (A, C, E) and Santa Cruz (B, D, F) anti-Kv1.3 subunit antibodies. Only weak or absent Kv1.3 staining was found in the soma of cerebellar output neurons in all three species (A, B). Some weak staining was observed in the surrounding neuropil. Scale bar=125μm.

## Discussion

The main objective of this study was to describe the cerebellar distribution of six Kv1 channel subunits (Kv1.1-1.6) in human, monkey and mouse and to determine whether our findings concur with the already known distribution of these proteins in rodents (Chung et al., 2001; Chung et al., 2005). This is the first study to directly examine inter-species (rodent to primate) and intra-species (human to non-human primate) differences in the distribution of these subunits in the cerebellum.

In general, the spatial patterning of the six proteins in the human and non-human primate cerebellum appeared strikingly similar. As expected, Kv1 channel subunits showed a very distinct cell-type specific pattern of expression in the cerebellum. Briefly, extensive overlapping of staining was seen for Kv1.1 and Kv1.2 subunits, which were predominantly found in the basket cell pinceau at the base of Purkinje cells, and strongest Kv1.3 and Kv1.5 immunoreactivities were present in somatodendric Purkinje cell areas. In general, human and monkeys showed similar patterns of expression as rodents for Kv1.1, Kv1.2, Kv1.3 and Kv1.5, but some marked differences for Kv1.4 and Kv1.6, with little or no Purkinje cell staining of Kv1.4 and Kv1.6. An additional novel finding of the study was intense stellate cell immunoreactivity, noted for the first time, but only in primate species.

Species differences in the spatial patterning of the six Kv1 channels have previously been described in other brain regions. For instance, in the gerbil hippocampus, Park et al., (2001) reported major differences in rodent distribution of the six Kv1 channel subtypes in comparison to the previous work in the rat and mouse. Recently, strain-specific hippocampal distribution of the Kv1 channel subtype-immunoreactivity has been reported in seizure sensitive and seizure resistant gerbils (Kim et al., 2007). Species-specific distribution of Kv1 channels has also been described in the rodent cerebellum. In gerbil, strong immunoreactivity for Kv1.4 has been observed in Purkinje cell bodies (Chung et al., 2005) in comparison with substantially lower staining in the rat (Chung et al., 2001). Kv1.2 immunoreactivity, albeit at low levels, was present in gerbil Purkinje cells bodies (Chung et al., 2005), but absent in the rat (Chung et al., 2001). It appears also that the distribution of Kv1 channels remains controversial even in the same species. For example, the presence of Kv1.2 staining is variable between studies in the rat alone (Chung et al., 2001; McNamara et al., 1993; McNamara et al., 1996; Sheng et al., 1994; Wang et al., 1994). Our study showed weak Kv1.2 staining in the Purkinje cells in mice and primates, and this is in agreement with some of those above and also *in situ* hybridization studies that have demonstrated that Purkinje cells express Kv1.2 mRNA (Kues and Wunder, 1992; Sheng et al., 1994; Tsaur et al., 1992). Such discrepancies may reflect issues of polyclonal Kv1.2 antibody specificity (McNamara et al., 1996), and the potential inherent differences between *in situ* hybridisation and immunocytochemistry in terms of differential translational efficiency of transcripts (Wang et al., 1994). Nonetheless, the current study (using the same antibodies as Chung et al., 2001; 2005) confirms findings previously made in rat and gerbil to those in a further rodent, and primates.

The main limitation of this study stems from the use of formalin-fixed paraffin-embedded tissues precluding use of other molecular techniques, such as *in situ* hybridization in order to confirm our findings. However, the Kv1 mRNA expression pattern obtained through *in situ* hybridization has not always been found to correlate well with the Kv1 immunostaining pattern (Veh et al., 1995). At present, we are unaware of any studies utilizing molecular techniques to corroborate the cerebellar distribution of the six Kv channels presented here, especially in primates. Specificity of immunoreactivity to the six Kv1 channels in the rodent brain has however already been confirmed by immunohistochemistry (Chung et al., 2000; Park et al., 2001; Guan et al., 2006) using in part antibodies used in our study (Alomone) in the gerbil and rat cerebellum (Chung et al., 2005 and Chung et al., 2001; respectively). The specificity of the Santa Cruz antibodies had not however been previously tested. Based on the manufacturers’ information provided for each of the commercially available antibody, it appears that each set of corresponding antibodies, e.g., Kv1.1 from Alomone vs Santa Cruz, was raised against a different epitope of the same protein (Table 1 & 2). The two sets of antibodies showed the same distribution patterns for Kv1.1, Kv1.2, Kv1.3, in particular, thus appearing to confirm specificity of these two sets of antibodies for these particular proteins. However, the differences seen in the distribution of Kv1.4, Kv1.5 and Kv1.6 channel subunits lead us to raise several other points. First of all, it emerged that differential distribution of Kv1.5 and Kv1.6 subunits between Alomone and Santa Cruz antibodies, especially in monkey and human tissue (Figs. 5C vs 5D; 5E vs 5F; 6C vs 6D; 6E vs 6F), casts a doubt on Santa Cruz antibodies giving a correct localization. Equally, absent staining of Kv1.4 (Fig. 4D, F) and Kv1.6 (Fig. 6D) could be due to poor Santa Cruz antibodies specificity. We have however no reason to believe that Kv1.5 Alomone antibody gave aberrant staining since it conferred staining seen in mouse (Fig. 5A, C, E). Moreover, Alomone Kv1.6 and Kv1.4 immunoreactivities showed consistently presence of stellate like cells evident not only within the cerebellar hemispheres (Fig. 4C, E; 6C, E), but also within the deep cerebellar nuclei (Fig. 10C, E; 12C, E). We thus speculate that the differences in the primate distribution of Kv1.5 most likely represent aberrant antibody staining rather than true species differences. However, Kv1.4 and Kv1.6 distributions appear to reflect real species differences, albeit this finding may be based only on our analysis of Kv1 Alomone set of antibodies.

**Figure 10.**
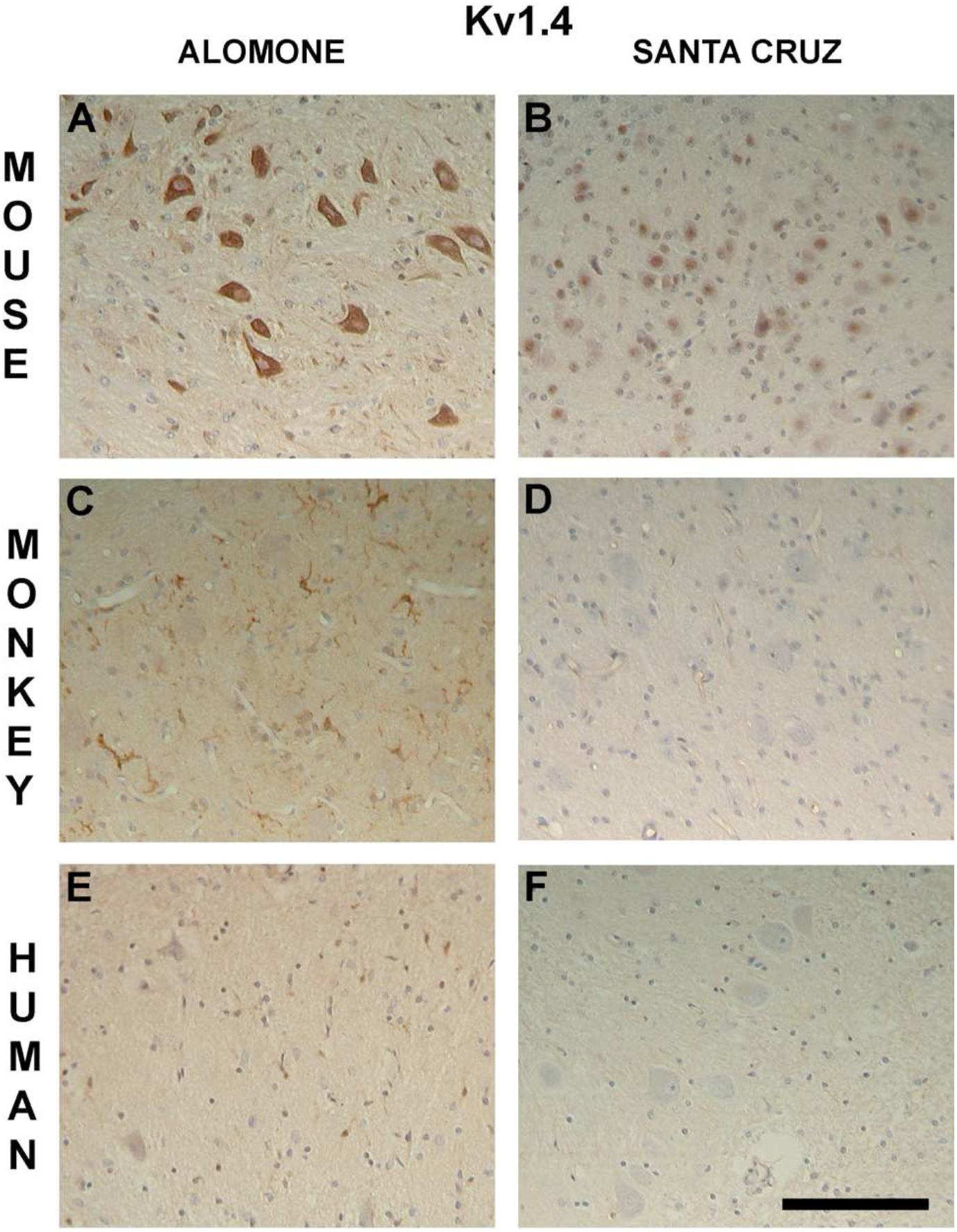
Cellular localizations of Kv1.4 subunit in the mouse (A, B); monkey (C, D) and human (E, F) cerebellar nuclei. Immunohistochemistry was obtained using Alomone (A, C, E) and Santa Cruz (B, D, F) anti-Kv1.4 subunit antibodies. In mouse, generally strong Kv1.4 staining was detected in the cell bodies of cerebellar output neurons, using both sets of antibodies (A, B). In contrast, in primates, staining was absent from the soma of cerebellar output neurons (C, D, E, F). In addition, in primates, using Alomone antibodies, relatively strong (C, E) immunoreactivity was found in cells, which appear to be stellate in form. Scale bar=125μm.

**Figure 11.**
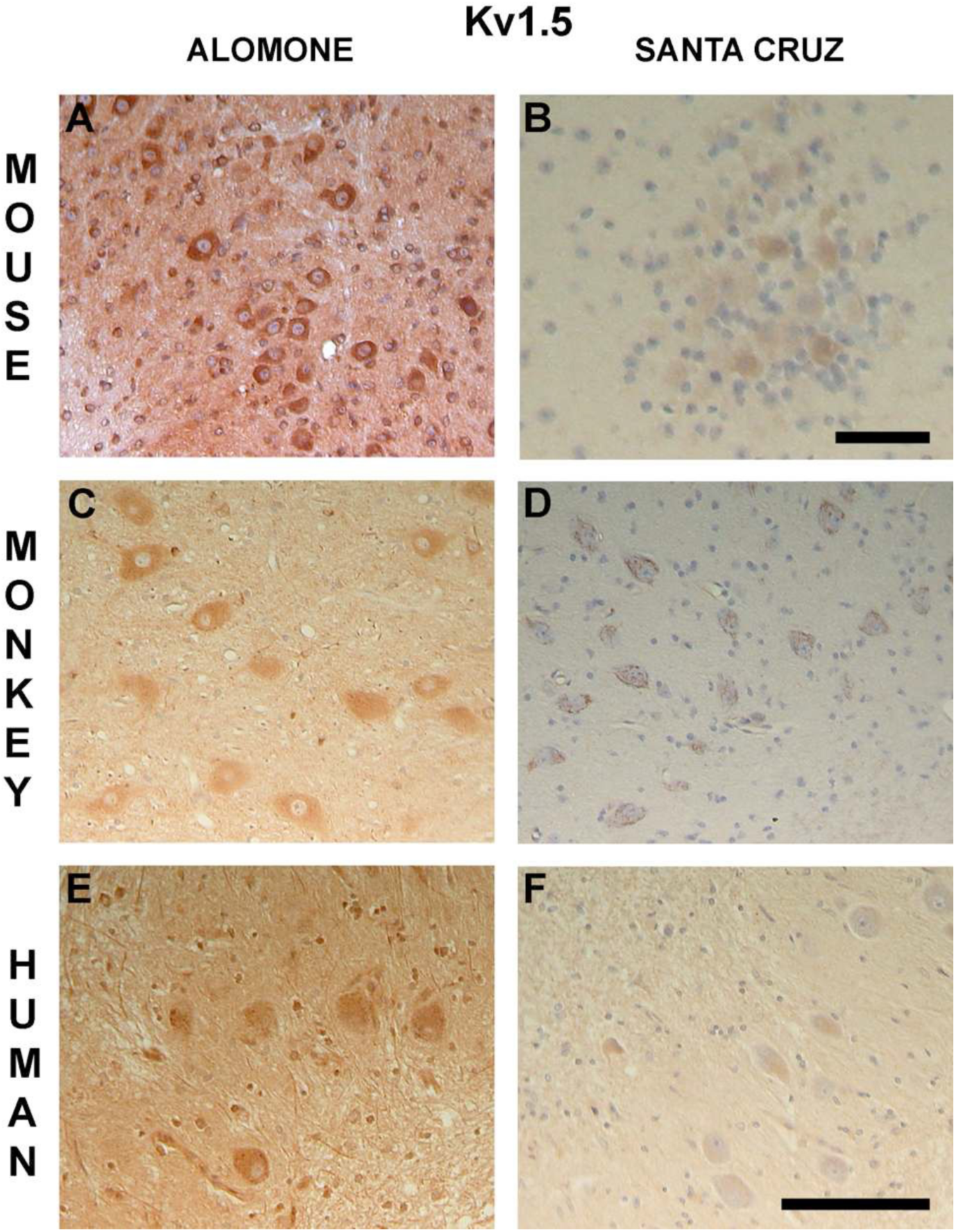
Cellular localizations of Kv1.5 subunit in the mouse (A, B); monkey (C, D) and human (E, F) cerebellar nuclei. Immunohistochemistry was obtained using Alomone (A, C, E) and Santa Cruz (B, D, F) anti-Kv1.5 subunit antibodies. Kv1.5 proteins were clearly detected in the cell bodies of cerebellar output neurons in all three species, using Alomone antibodies (A, C, E). Similar, although weaker, cell soma staining pattern was observed in mouse (B) and monkey (D), and the weakest in human (F), using Santa Cruz antibodies. Some staining in the surrounding neuropil was also observed. Scale bar=100μm in (B), otherwise scale bar=125μm.

**Figure 12.**
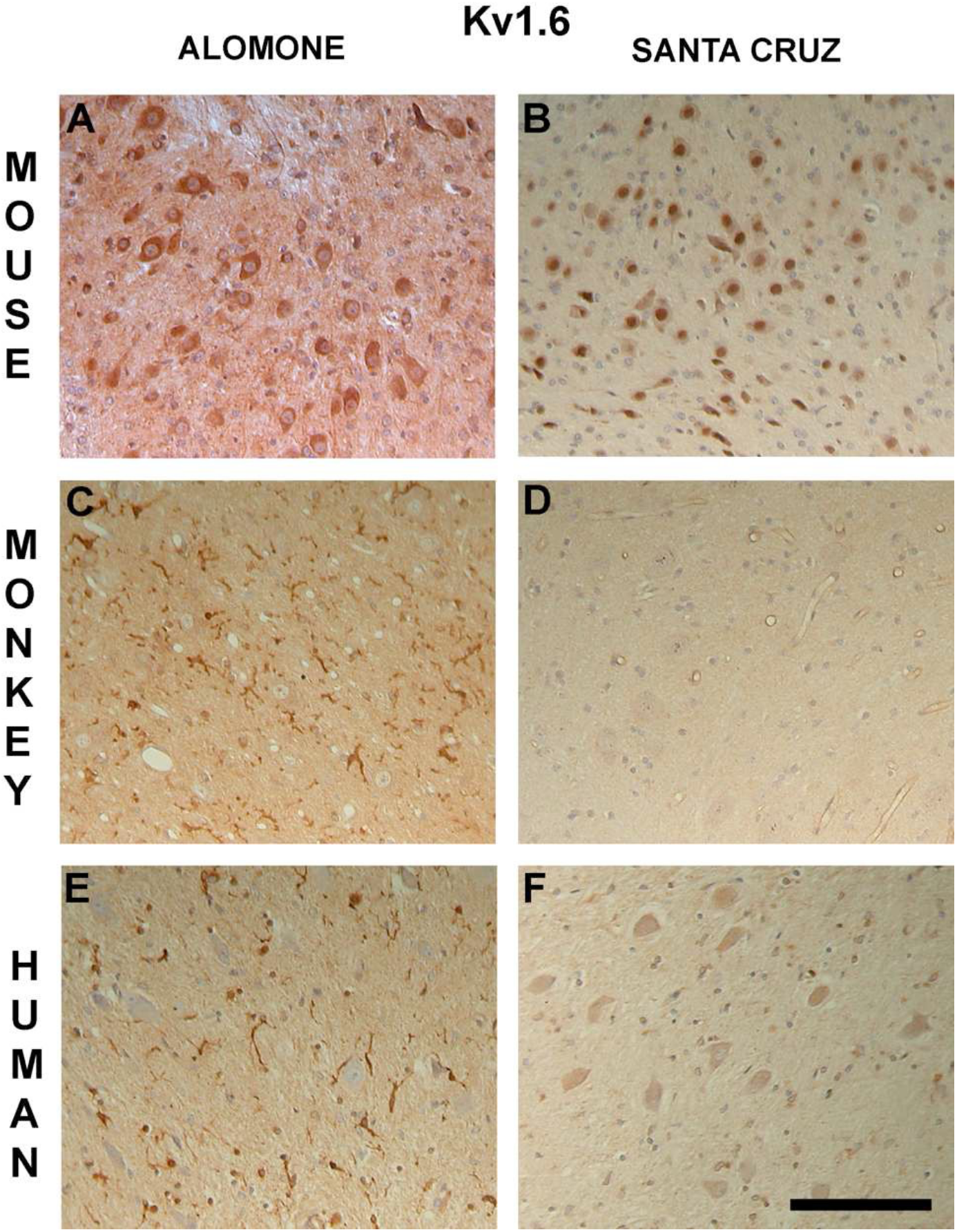
Cellular localizations of Kv1.6 subunit in the mouse (A, B); monkey (C, D) and human (E, F) cerebellar nuclei. Immunohistochemistry was obtained using Alomone (A, C, E) and Santa Cruz (B, D, F) anti-Kv1.6 subunit antibodies. Kv1.6 immunoreactivity was predominantly found in the cell bodies of cerebellar output neurons in mouse, using both antibodies (A, B). However, only weak (C, D, F) or absent (E) signal was found in cell bodies in primates. In addition, in primates, using Alomone antibodies, we found relatively strong immunoreactivity for what appear to be stellate in form cells (C, E). Scale bar=125μm.

The present study describes only the distribution of Kv1 subunits. The functional implications of our findings can only be projected. The pattern of expression of different Kv1 subunits may correspond to their pattern of biophysical properties. Except for Kv1.4, Kv1 channel subunits form slowly inactivating currents when expressed as homomeric channels in heterologous systems. Kv1.4 channel subunit homomers form a rapidly inactivating A-type current (Pongs, 1992). However, our knowledge about specific roles of native channels or the subunit composition is still limited. Usually, the availability of Kv channel mutants is of great help in identifying native Kv channel properties (Pongs, 1992). However, comparable observations with *Drosophila* (Lichtinghagen et al., 1990) and mice (Smart et al., 1998) in which the functional expression of certain Kv channels subunits were knocked out demonstrated that phenotypes associated with null mutants were not readily apparent. Few known human Kv channel genes have been correlated with disease phenotypes (Pongs, 1999). In humans, mutations in a gene *KCNA1* encoding Kv1.1 channel have been linked to episodic ataxia type-1 (EA-1), an autosomal dominant neurological disorder characterized by cerebellar ataxia (Browne et al., 1994). However, even mutations in the *KCNA1* lead only to a reduction rather than loss of function (Adelman et al., 1995). It is thus not always possible to correlate loss of function of a certain Kv channel subunit phenotypic changes in neuronal excitability, this being probably due to the molecular complexity of Kv channels (Pongs, 1999).

This study has described the cerebellar distribution of six Kv1 channels subunits in three more species than previously studied and adds to the evidence of species-specific cellular expression of certain Kv channels. Such studies may prove important in the future if inferences to human disease pathways or therapy are to be made from data gathered solely in rodent models.

## Acknowledgements

We thank Alex Brown, Carole Nickols and Chris Evagora for technical assistance. Part of this work was presented at the 109^th^ Meeting of the British Neuropathological Society, January 9-11, 2008, London, UK.

## Literature Cited

Adelman JP, Bond CT, Pessia M, Maylie J (1995) Episodic ataxia results from voltage-dependent potassium channels with altered functions. Neuron 15:1449–1454.

Browne DL, Gancher ST, Nutt JG, Brunt ER, Smith EA, Kramer P, Litt M (1994) Episodic ataxia/myokymia syndrome is associated with point mutations in the human potassium channel gene, KCNA1. Nat Genet 8:136–140.

Chung YH, Shin CM, Kim MJ, Cha CI (2000) Immunohistochemical study on the distribution of six members of the Kv1 channel subunits in the rat basal ganglia. Brain Res 875:164–170.

Chung YH, Shin C, Kim MJ, Lee BK, Cha CI (2001) Immunohistochemical study on the distribution of six members of the Kv1 channel subunits in the rat cerebellum. Brain Res 895:173–177.

Chung YH, Joo KM, Nam RH, Kim YS, Lee WB, Cha CI (2005) Immunohistochemical study on the distribution of the voltage-gated potassium channels in the gerbil cerebellum. Neurosci Lett 374:58–62.

Guan D, Lee JC, Tkatch T, Surmeier DJ, Armstrong WE, Foehring RC (2006) Expression and biophysical properties of Kv1 channels in supragranular neocortical pyramidal neurones. J Physiol 571:371–389.

Gutman GA, Chandy KG, Grissmer S, Lazdunski M, McKinnon D, Pardo LA, Robertson GA, Rudy B, Sanguinetti MC, Stuhmer W, Wang X (2005) International Union of Pharmacology. LIII. Nomenclature and molecular relationships of voltage-gated potassium channels. Pharmacol Rev 57:473–508.

Hille B (2001) Ion channels of excitable membranes. Sunderland, MA: Sinauer.

Kim DS, Kim JE, Kwak SE, Won MH, Kang TC (2007) Seizure activity affects neuroglial Kv1 channel immunoreactivities in the gerbil hippocampus. Brain Res 1151:172–187.

Kues WA, Wunder F (1992) Heterogeneous Expression Patterns of Mammalian Potassium Channel Genes in Developing and Adult Rat Brain. Eur J Neurosci 4:1296–1308.

Lichtinghagen R, Stocker M, Wittka R, Boheim G, Stuhmer W, Ferrus A, Pongs O (1990) Molecular basis of altered excitability in Shaker mutants of Drosophila melanogaster. Embo J 9:4399–4407.

McNamara NM, Muniz ZM, Wilkin GP, Dolly JO (1993) Prominent location of a K+ channel containing the alpha subunit Kv 1.2 in the basket cell nerve terminals of rat cerebellum. Neuroscience 57:1039–1045.

McNamara NM, Averill S, Wilkin GP, Dolly JO, Priestley JV (1996) Ultrastructural localization of a voltage-gated K+ channel alpha subunit (KV 1.2) in the rat cerebellum. Eur J Neurosci 8:688–699.

Park KH, Chung YH, Shin C, Kim MJ, Lee BK, Cho SS, Cha CI (2001) Immunohistochemical study on the distribution of the voltage-gated potassium channels in the gerbil hippocampus. Neurosci Lett 298:29–32.

Pongs O (1992) Molecular biology of voltage-dependent potassium channels. Physiol Rev 72: S69–88.

Pongs O (1999) Voltage-gated potassium channels: from hyperexcitability to excitement. FEBS Lett 452:31–35.

Sheng M, Tsaur ML, Jan YN, Jan LY (1992) Subcellular segregation of two A-type K+ channel proteins in rat central neurons. Neuron 9:271–284.

Sheng M, Liao YJ, Jan YN, Jan LY (1993) Presynaptic A-current based on heteromultimeric K+ channels detected in vivo. Nature 365:72–75.

Sheng M, Tsaur ML, Jan YN, Jan LY (1994) Contrasting subcellular localization of the Kv1.2 K+ channel subunit in different neurons of rat brain. J Neurosci 14:2408–2417.

Smart SL, Lopantsev V, Zhang CL, Robbins CA, Wang H, Chiu SY, Schwartzkroin PA, Messing A, Tempel BL (1998) Deletion of the K(V)1.1 potassium channel causes epilepsy in mice. Neuron 20:809–819.

Tsaur ML, Sheng M, Lowenstein DH, Jan YN, Jan LY (1992) Differential expression of K+ channel mRNAs in the rat brain and down-regulation in the hippocampus following seizures. Neuron 8:1055–1067.

Veh RW, Lichtinghagen R, Sewing S, Wunder F, Grumbach IM, Pongs O (1995) Immunohistochemical localization of five members of the Kv1 channel subunits: contrasting subcellular locations and neuron-specific co-localizations in rat brain. Eur J Neurosci 7:2189–2205.

Wang H, Kunkel DD, Schwartzkroin PA, Tempel BL (1994) Localization of Kv1.1 and Kv1.2, two K channel proteins, to synaptic terminals, somata, and dendrites in the mouse brain. J Neurosci 14:4588–4599.

Wang H, Kunkel DD, Martin TM, Schwartzkroin PA, Tempel BL (1993) Heteromultimeric K+ channels in terminal and juxtaparanodal regions of neurons. Nature 365:75–79.

